# Understanding properties of the master effector of phage shock operon in *Mycobacterium tuberculosis* via bioinformatics approach

**DOI:** 10.1101/050047

**Authors:** Zarrin Basharat, Azra Yasmin

## Abstract

The phage shock protein (Psp) is a part of the Psp operon, which assists in safeguarding the survival of bacterium in stress and shields the cell against proton motif force challenge. It is strongly induced by bacterium allied phages, improperly localized mutant porins and various other stresses. Master effector of the operon, PspA has been modeled and simulated, illustrating how it undergoes significant conformational transition at the far end in *Mycobacterium tuberculosis*. Association of this key protein of the operon influences action of Psp system on the whole. We are further working on the impact of phosphorylation perturbation and changes in the structure of PspA during complex formation with other moieties of interest.

---

We hereby report *ab initio* structure model of the *Mycobacterium tuberculosis* phage shock protein A (PspA). PspA is the central constituent of bacterial stress response machinery, encoded by phage shock operon (Huvet *et al.*, 2010). PspA, regulates not only it’s own transcription but that of the whole operon as well (Elderkin *et al.*, 2005; Male *et al.*, 2014). PspA is inferred to be a dual-function protein (Guegwen *et al.*, 2009; Jovanovic *et al.*, 2014) and localized amid cytoplasmic and inner membrane interface of the bacterium (Engl *et al.*, 2009). It is responsible for maintaining the cell membrane integrity along with restoration of the proton motive force (Male *et al.*, 2014; Engl *et al.*, 2009; Wan *et al.*, 2015).

*Mycobacterium tuberculosis* is a pathogenic bacterium responsible for causing disease in humans and veterinary species (Sakamoto, 2012). A lot of work has been carried out on *Mycobacterium* phages for therapeutic purpose. However, to the best of our knowledge, no report of the phage shock protein analysis for this pathogen exists at the moment. We retrieved PspA protein sequence of *Mycobacterium tuberculosis* from the Uniprot database with Accession number: R4M912 and analyzed the sequence and the structure using computational tools. Although, PspA belongs to the highy conserved PspA/IM30 family but *Mycobacterium tuberculosis* PspA shared a very low sequence homology with *Escherichia coli* PspA, revealed using Clustal Omega (Fig. 1) with seeded guide trees and HMM profile-profile technique for alignment generation at the backend (Sievers *et al.*, 2011). The secondary structure analyzed by PROMOTIF tool (Hutchison and Thornton, 1996) revealed that the protein consisted of 9 helices, 6 helix-helix interacs, 14 β-turns and 3 □-turns (Fig. 2A).

**Figure 1.**
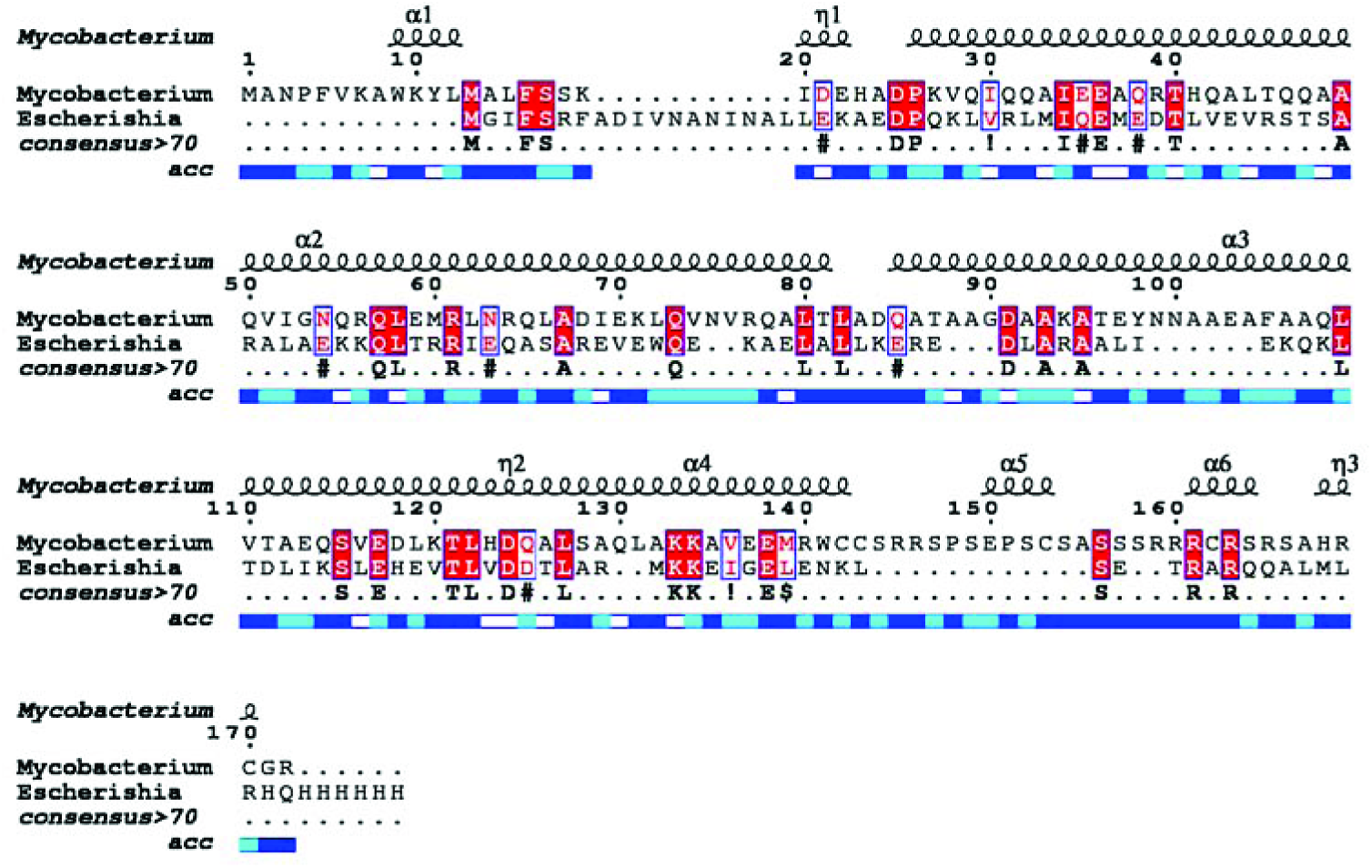
Multiple sequence alignment of the *Escherichia coli* and *Mycobacterium tuberculosis* PspA. Conserved residues are shown in red color. Helices are denoted by squiggles at the top of the alignment. Solvent accessibility is depicted by a bar below the sequence (blue = accessible, cyan = intermediate, white = buried).

**Figure 2.**
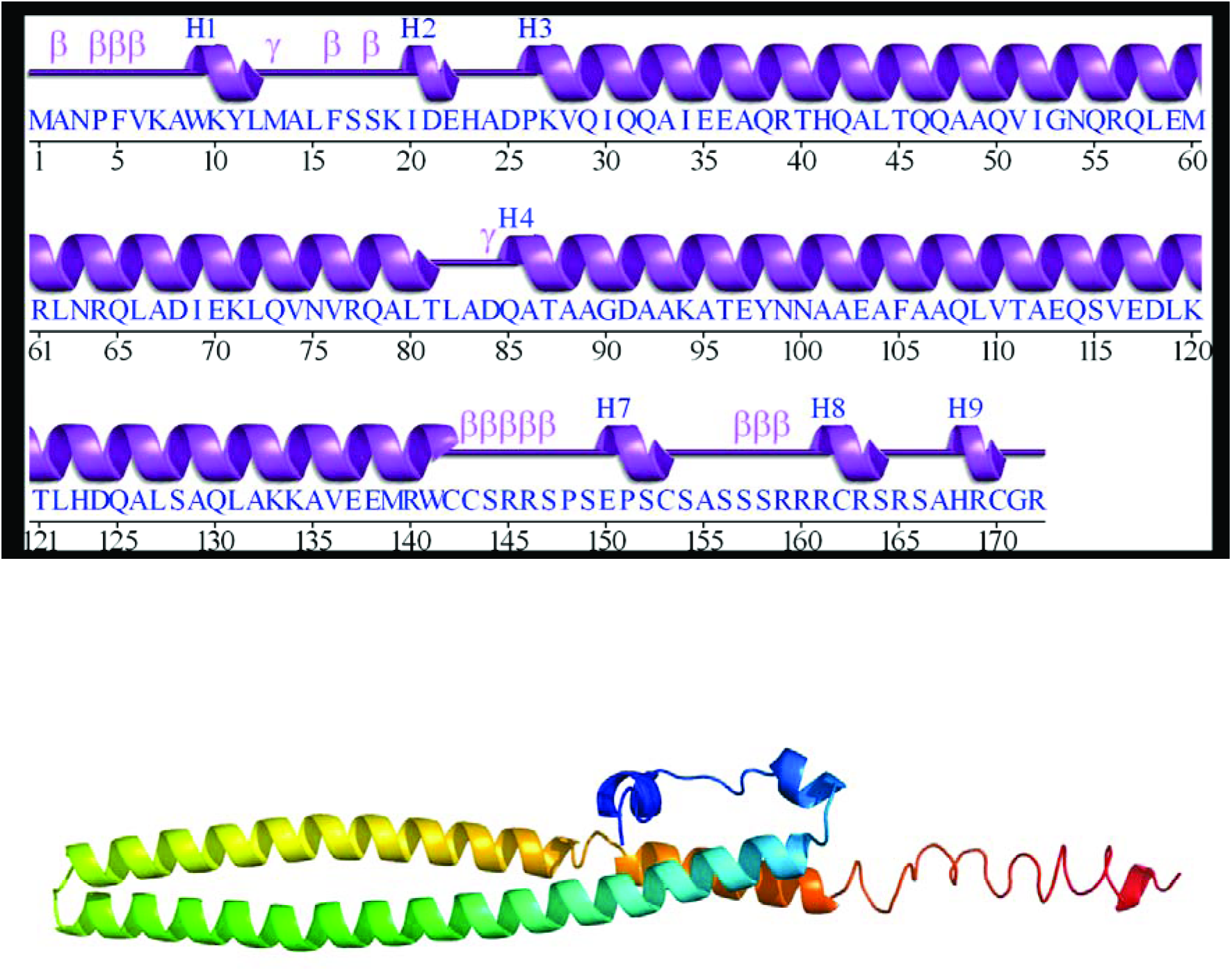
(A) Secondary structure *of Mycobacterium tuberculosis* PspA (helices labelled H1, H2…±H9). **P** depicts beta turn and is for gamma turn. (B) 3D structure *of Mycobacterium tuberculosis* PspA.

Due to low homology with experimentally determined structures available in the RCSB Protein databank, 3D structure (Fig. 2B) was modeled by I-TASSER (Roy *et al.*, 2010; Yang *et al.*, 2015) using *Escherichia coli* PspA as a template (PDB ID: 4WHE). The C-score based on the significance of template alignment threading and simulations of the structure assembly convergence parameters of the model chosen for analysis was −1.29 (lies between ideal range of −5 to 2), indicating good quality model. Estimated root mean square deviation of the predicted model from the *Escherichia coli* model was 7.8±4.4Å. The constructed structure resembled a helical bundle with important binding site residues predicted to occur at position 66, 69, 70, 73, 92, 96, 99, 100, 102 and 103. Despite low sequence conservation, the structure is however, well conserved due to the underlying fact that the protein folds remain conserved in similar function proteins. PspA structure in *Escherichia coli* is known to self-assemble into ring (Standar *et al.*, 2008) or striated and indented rod-shaped complexes (Male *et al.*, 2014) based on electron microscopy and helical rods based on X-ray crystallography analysis (Osadnik *et al.*, 2015). PspA homologue LiaH in *Bacillus subtilis* (Wolf *et al.*, 2008) and holins of bacteriophage lambda (Savva *et al.*, 2010) have also been reported to self-assemble to rod-like structures from ring shaped protein complexes. The *Mycobacterium tuberculosis* PspA is also rod shaped and it is implied that these rod-like structures could form a support framework and aid in the maintenance of membrane integrity during phage shock response (Male *et al.*, 2014).

CABS-flex procedure based on the well-established coarse-grained CABS protein model (Fraga *et al.*, 2014) was employed for the fast simulation of near-native dynamics of PspA. CABS is a computationally efficient alternative to classical all-atom molecular dynamics (Jamroz *et al.*, 2013). The 3D modeled structure was input and used as a starting point for the all-atom, explicit water, 10-nanosecond dynamic simulation. Analysis was carried out at the backend and automatically analyzed trajectory (Fig. 3) was obtained to study the dynamic behaviour of protein. A set of eight (all-atom) protein model sets were obtained with global distance test score ranging from 0.6-0.7. Most dominant structural fluctuations appeared at the last beta turn region including 8^th^ and 9^th^ helix. Relative propensity of protein residues to deviate from an average dynamics structure increased substantially at the ending helices with fluctuation increasing from 100 Å at residue 160 to to >600 Å at residue 172. Understanding of flexibility of PspA can be of aid in research areas as molecular evolution (Boehr *et al.*, 2009).

**Figure 3.**
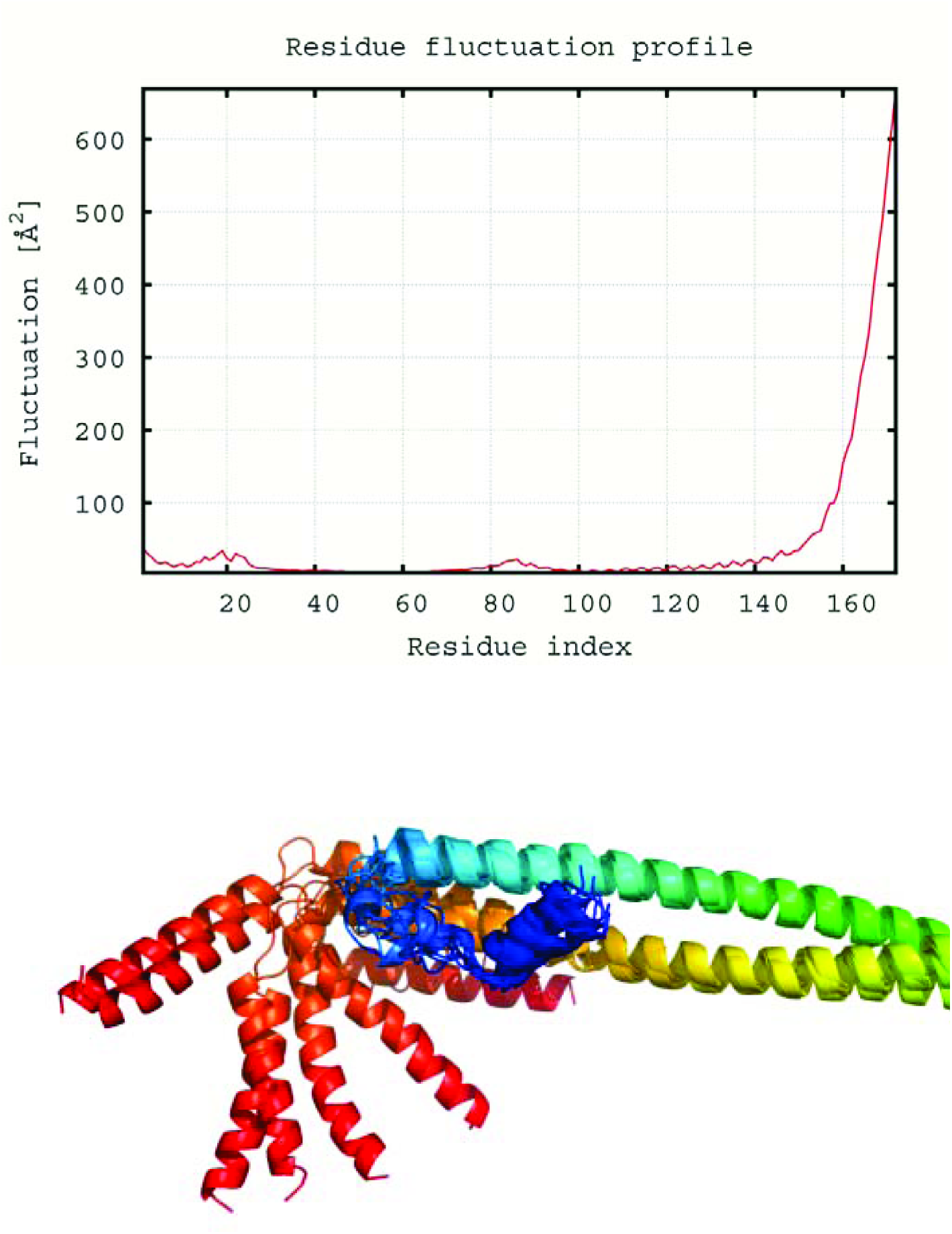
(A) Structural flexibility profile of simulated *Mycobacterium tuberculosis* PspA with fluctuations for individual protein residues shown via red line. The output is based on all-atom model via trajectory clustering. (B) Refinement of the model and superpositioning (Provided 3D model as base) is centred on maximum likelihood superposition method of THESEUS (Theobald and Wuttke, 2006).

Phage infection has also been demonstrated to induce substantial fluctuations in host protein phosphorylation (Rieul *et al.*, 1987; Russel and Model, 2006) and this was suggestive of PspA potential for phosphorylation as well. NetPhos Bac 1.0 (Miller *et al.*, 2009) was used for prediction of possible phosphorylation residues. Seven serine residues (S144, S149, S156, S157, S158, S164, S166) were predicted to have phosphorylation potential based on neural network approach. However, none of these exhibited a knack to occur on predicted binding residues and hence, their exact role in PspA function of *Mycobacterium tuberculosis* yet remains to be elucidated.

Our findings pave way for further experimental studies and are of aid in understanding the *Mycobacterium tuberculosis* PspA response to the extracytoplasmic stresses that may damage the cytoplasmic membrane. We have used computational approach for the prediction of 3D structure of this protein but to furthur understand the function of the rod-like structure of *Mycobacterium tuberculosis* PspA, additional studies are required which can confirm and enhance the reported information along with elucidation of in depth biological function and interactions of *Mycobacterium tuberculosis* PspA.

## Acknowledgements

The authors are thankful to Alexandra Elbakyan for providing literature support.

